# CanVAS: A Harmonized and Imputed Canine Variant Atlas

**DOI:** 10.64898/2026.04.13.718238

**Authors:** David M. Brundage

## Abstract

The domestic dog (*Canis lupus familiaris*) is a powerful model for genetic studies of complex disease, but canine genotype data are distributed across independent studies using incompatible genotyping platforms, genome builds, strand conventions, and allele coding schemes. Here we present CanVAS, a quality-controlled, harmonized, and imputed canine genotype resource integrating 15 publicly available datasets into a single analysis-ready PLINK file set on the CanFam4 (UU_Cfam_GSD_1.0) reference assembly. The typed backbone contains 15,451 dogs from over 375 breeds, village dog populations, dingoes, wolves, and coyotes, genotyped across 77,215 shared SNPs. Imputation against the Dog10K whole-genome sequencing reference panel (1,929 dogs) using Beagle 5.4 expanded the resource to 9.7 million variants (DR2 ≥ 0.3, MAF ≥ 0.01), including approximately 3 million rare variants (MAF *<* 0.05). We describe the complete harmonization pipeline and validate the resource through population structure analysis and genome-wide runs-of-homozygosity analysis, recovering known breed-level differences in genomic inbreeding.

## 1 Background & Summary

The domestic dog offers unique advantages for genetic mapping of complex traits and diseases. Approximately 450 recognized breeds, each founded from small numbers of individuals within the past few hundred years, create natural genetic isolates with extended linkage disequilibrium, high trait heritability, and reduced allelic heterogeneity relative to human populations [Ostrander, 2012, Lindblad-Toh et al., 2005]. These properties have enabled genome-wide association studies (GWAS) to identify risk loci for complex diseases including osteosarcoma [Karlsson et al., 2013], histiocytic sarcoma [Hédan et al., 2021], mast cell tumour [Biasoli et al., 2019], and numerous Mendelian disorders, often with substantially fewer samples than required in human studies.

Over the past decade, canine GWAS datasets have accumulated across independent research groups worldwide, genotyped primarily on the Illumina CanineHD BeadChip (∼170,000 SNPs) and the Thermo Fisher Axiom Canine array (∼710,000–1.1M SNPs). Many of these datasets have been deposited in public repositories under open licenses. However, cross-cohort integration has been limited by several technical barriers: incompatible SNP identifier formats (Illumina probe names vs. Axiom probe names vs. positional IDs), mixed genome builds (CanFam2, CanFam3.1, and now CanFam4), divergent strand conventions (Illumina TOP/BOT vs. forward strand), and inconsistent allele coding schemes (ACGT vs. numeric). As a result, each research group typically analyzes only its own data, leaving substantial statistical power unrealized.

The recent release of the Dog10K whole-genome sequencing reference panel [Dog10K, 2024]—comprising 1,929 canids from 321 breeds—has created an unprecedented opportunity for genotype imputation of SNP array data to whole-genome density. However, exploiting this panel requires a harmonized, multi-cohort target dataset on a consistent coordinate system, which has not previously existed.

Here we address these gaps by presenting CanVAS, a harmonized and imputed genotype resource integrating 15 publicly available canine datasets into a single PLINK file set on the CanFam4 reference assembly. The resource encompasses 15,451 dogs spanning over 375 breeds, Australian dingoes, Chinese indigenous dogs, village dogs from five continents, wolves, and coyotes. The typed backbone of 77,215 shared SNPs has been imputed to 9.7 million variants using the Dog10K reference panel, transforming a common-variant array scaffold into a dataset that captures the full allele frequency spectrum.

We describe the complete harmonization pipeline, provide all scripts for reproducibility, and validate the resource through population structure analysis, imputation quality assessment, and a genome-wide runs-of-homozygosity (ROH) analysis that recovers known breed-level inbreeding patterns. The intended uses of this resource include population structure analysis, cross-breed association studies and meta-analyses, polygenic risk score portability evaluation, imputation benchmarking, and ROH-based inbreeding and selection sweep mapping across the purebred-to-village dog continuum.

## 2 Methods

### 2.1 Source datasets

We identified 15 publicly available canine genotype datasets deposited in Dryad, Zenodo, or accessible through institutional data commons (Table 1). Selection criteria included: (1) genotyping on the Illumina CanineHD, Thermo Fisher Axiom, or whole-genome sequencing platforms; (2) availability of individual-level genotype data in PLINK or VCF format; and (3) deposition under CC0 or compatible open-access terms. The largest cohorts are Hayward et al. 2016 (*n* = 4,339; Illumina CanineHD) [Hayward et al., 2016,b], the Golden Retriever Lifetime Study (GRLS; *n* = 3,197; Axiom Canine HD) [GRLS, 2024], and Spatola et al. (*n* = 1,903) [Spatola et al., 2022]. Together, the 15 cohorts encompass 15,451 dogs from over 375 breeds.

**Table 1:** Source datasets integrated into CanVAS.

| # | Dataset | Dogs | Typed SNPs | Platform | Original Build | Citation |
| --- | --- | --- | --- | --- | --- | --- |
| 1 | Hayward 2016 | 4,339 | 160,723 | Illumina HD | CanFam3.1 | Hayward et al. [2016,b] |
| 2 | GRLS | 3,197 | 802,053 | Axiom | CanFam3.1 | GRLS [2024] |
| 3 | Spatola | 1,903 | 129,497 | Illumina HD | CanFam3.1 | Spatola et al. [2023, 2022] |
| 4 | Shannon | 1,180 | 185,805 | Illumina custom | CanFam3.1 | Shannon et al. [2015, 2016b] |
| 5 | Parker 2017 | 1,031 | 148,704 | Illumina HD | CanFam3.1 | Parker et al. [2017,b] |
| 6 | Wiener 2017 | 726 | 115,993 | Illumina HD | CanFam2→3.1 | Wiener et al. [2017, 2018] |
| 7 | Hédan 2021 | 589 | 117,030<sup>a</sup> | Mixed<sup>b</sup> | CanFam3.1 | Hédan et al. [2021, 2023] |
| 8 | Momozawa 2020 | 543 | 167,712 | Illumina HD | CanFam2→3.1 | Momozawa et al. [2020,b] |
| 9 | Cairns 2023 | 531 | 192,559 | Axiom HD | CanFam3.1 | Cairns et al. [2023,b] |
| 10 | HaywardDeaf 2020 | 503 | 181,252 | Illumina 220K | None→3.1 | Hayward et al. [2020] |
| 11 | Chen/Yang 2019 | 325 | 131,763 | Illumina HD | None→3.1 | Chen et al. [2019] |
| 12 | Biasoli 2019 | 279 | 163,361 | Illumina HD | CanFam2→3.1 | Biasoli et al. [2019,b] |
| 13 | JenkinsWGS 2021 | 180 | 633,325 | WGS extract | CanFam3.1 | Jenkins et al. [2021] |
| 14 | Bianchi 2020 | 115 | 169,445 | Illumina HD | CanFam3.1 | Bianchi et al. [2015] |
| 15 | Bespalova | 8 | 172,115 | Illumina HD | None→3.1 | Bespalova et al. [2025,b] |
| **Total** |  | **15,451** |  |  |  |  |

<sup>a</sup>Illumina subset; Axiom subset contains 582,165 variants.
<sup>b</sup>457 Illumina CanineHD + 134 Axiom 1.1M, split and harmonized independently.

**Table 2:** Allele frequency divergence at known trait loci between breed groups with opposing phenotypes.

| Locus | Phenotype contrast | Typed Δfreq | Imputed Δfreq |
| --- | --- | --- | --- |
| IGF1 (chr15) | Large vs. small breeds | 0.211 | 0.453 |
| LCORL (chr3) | Large vs. small breeds | 0.290 | 0.448 |
| FGF5 (chr32) | Long vs. short coat | 0.365 | 0.401 |
| MC1R (chr5) | Yellow/red vs. dark coat | 0.337 | 0.459 |

### 2.2 Per-cohort quality control

Each cohort was processed independently prior to cross-cohort merging using PLINK v1.9 [Chang et al., 2015] with the `–dog flag`. For datasets where universally missing variants inflated per-sample missingness (e.g., Momozawa, overall genotyping rate 93.1%), variant missingness filtering (`–geno 0.02`) was applied before sample missingness filtering (`–mind 0.05`) to avoid spurious sample exclusion. SNPs with *>*2% missing genotypes were removed, followed by dogs with *>*5% missing genotypes. Hardy–Weinberg equilibrium filtering (*p <* 1 × 10^*−*6^) was applied only to single-breed cohorts (GRLS); for multi-breed cohorts, HWE testing was omitted because the Wahlund effect generates systematic departures that are biologically expected rather than indicative of genotyping error. This decision was empirically validated on the Parker dataset, where HWE filtering would have removed 97% of variants.

### 2.3 Genome build harmonization

All cohorts were initially harmonized to CanFam3.1 [Hoeppner et al., 2014]. Three classes of build issues were resolved:

#### CanFam2 coordinates (Wiener, Biasoli, Momozawa)

Three datasets contained positions on the obsolete CanFam2 build. Rather than using chain-file liftover, we exploited the fact that all three used Illumina CanineHD probe identifiers shared with the Hayward et al. dataset on CanFam3.1. Probe IDs served as stable cross-build keys: for each variant, the CanFam3.1 position was looked up from the Hayward BIM file by exact probe ID match, with fallback to prefix matching (splitting on `_rs` suffix) for truncated probe IDs. This approach ensured exact probe-level correspondence without chain-file approximations.

#### Missing coordinates (Yang/Chen, Hayward deafness)

Two datasets contained no genomic coordinates (chr=0, pos=0), retaining only probe identifiers. The same probe-ID-based mapping was applied using the Hayward BIM as reference.

#### CanFam3.1 to CanFam4 liftover

The harmonized dataset was lifted from CanFam3.1 to CanFam4 (UU_Cfam_GSD_1.0) [Wang et al., 2021] using per-chromosome VCF conversion. Chromosomes 27 and 32 required coordinate reversal, consistent with the documented orientation change [Wang et al., 2021, Field et al., 2020]. Of 77,344 CanFam3.1 backbone variants, 77,215 mapped successfully (129 unmappable).

### 2.4 Strand harmonization

Strand convention was determined empirically for each cohort via allele concordance analysis against the GRLS dataset chosen as the strand reference because it is the largest single-platform cohort (Axiom, forward strand). For each shared variant, allele pairs were classified as same-strand, complement (indicating Illumina TOP/BOT convention), monomorphic/zero-allele, or mismatch. Four datasets exhibited the characteristic ∼ 50/50 same-strand/complement split diagnostic of TOP/BOT convention; flip lists were generated and applied. Ambiguous A/T and C/G SNPs were excluded at merge time.

### 2.5 Allele recoding

The Wiener and Spatola dataset used Illumina GenomeStudio’s numeric 1/2 convention. A lookup table from the Hayward BIM mapped each probe to nucleotide alleles; recoding was applied at the PED genotype level and validated by tracing individual genotypes at five loci across ten dogs. The Cairns dataset used 1/2/3/4 coding, mapped empirically against GRLS as 1=A, 2=C, 3=G, 4=T.

### 2.6 Merging and shared backbone

All SNP IDs were standardized to `chr:pos` format. Cohorts were merged in two branches (Branch A: Parker + Hayward + GRLS + Wiener; Branch B: remaining cohorts), then combined. Triallelic variants identified by PLINK’s missnp procedure were excluded. The intersection of variants present in ≥90% of cohorts with per-variant call rate ≥90% defined the shared backbone: 77,344 variants on CanFam3.1, 77,215 after liftover to CanFam4.

### 2.7 Sample deduplication

Identity-by-descent analysis on the shared backbone (`–genome –min 0.90`) identified duplicate samples across cohorts. Pairs with 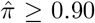 were resolved by retaining the sample from the higher-priority cohort. An additional 58 samples overlapping with the Dog10K reference panel were removed. After deduplication 14,478 dogs were remaining for analysis.

### 2.8 Breed annotation

A total of 439 raw breed labels were harmonized to 386 canonical breed names using a curated mapping resolving inconsistencies in case, delimiters, abbreviations, and regional naming. Breed groups used in figures were assigned using AKC classifications supplemented by keyword matching; these annotations are not included in the deposited metadata but can be derived from the breed labels using the provided scripts.

### 2.9 Imputation

Imputation was performed using Beagle 5.4 [Browning et al., 2018] with the Dog10K reference panel (1,929 dogs, 321 breeds, CanFam4) [Dog10K, 2024]. Per-chromosome jobs were run on the University of Wisconsin– Madison CHTC HTCondor cluster inside Apptainer containers. Post-imputation filtering retained variants with DR2 ≥ 0.3 and MAF ≥ 0.01.

### 2.10 Ethics statement

This study used only publicly available genotype datasets deposited in open-access repositories (Dryad, Zenodo, and institutional data commons) under CC0 or compatible open-access licenses. No new biological samples were collected and no animals were handled as part of this work. Ethics approval for the original data collection was obtained by the respective study investigators as described in the primary publications.

## 3 Data Records

The CanVAS resource is deposited on Zenodo Brundage [2025] and GitHub (https://github.com/Brundage-VAIL/CanVAS) and consists of:

1. **Typed backbone** (`CanVAS_shared_90pct_v4_canfam4.bed/bim/fam`): 77,215 variants × 15,451 dogs on CanFam4. Available on GitHub.
2. **Imputed dataset** (`CanVAS_v4_imputed_maf0.01.bed/bim/fam`): 9,667,790 variants (DR2 ≥ 0.3, MAF ≥ 0.01) on CanFam4. Available on Zenodo.
3. **Metadata** (`CanVAS_v4_metadata.tsv`): Per-dog cohort assignment, breed (386 canonical labels), and phenotype annotations. Available on Zenodo.
4. **QC summary** (`qc_chr_summary.tsv`): Per-chromosome imputation quality statistics. Available on Zenodo.
5. **Pipeline code**: All harmonization, imputation, and analysis scripts available on GitHub under MIT license.

Per-chromosome imputation files in VCF format retaining DR2 annotations can be regenerated from the imputed PLINK files using the provided scripts.

The metadata file contains the following fields: `FID` (PLINK family identifier), `IID` (PLINK individual identifier, unique per dog), `Cohort` (source dataset name, corresponding to Table 1), `Breed` (harmonized canonical breed label from 386 standardized names), `BreedDetail` (additional breed or population annotation where available), and `Phenotype` (semicolon-delimited phenotype labels from the source dataset, where available). Fields may be empty where annotations were not provided in the source data.

## 4 Technical Validation

### 4.1 Allele concordance

Strand and allele consistency was validated for every cohort against the GRLS reference. Concordance exceeded 99% for all forward-strand datasets after harmonization. TOP/BOT datasets showed the expected ∼50/50 same-strand/complement distribution, with *<*0.1% true mismatches after strand correction. The elevated apparent mismatch in the Bianchi cohort (21.7%) was diagnosed as monomorphic sites in this small single-breed cohort (115 Giant Schnauzers); only 2 of 139,142 shared sites were true mismatches.

### 4.2 Population structure

Principal component analysis on the LD-pruned backbone (PLINK `–indep-pairwise 50 5 0.2, 20 eigenvectors`) revealed strong breed-level structure, with the top four PCs explaining 25.2%, 14.2%, 8.4%, and 7.6% of variance. PC1 separates Golden Retrievers; PC2 distinguishes Boxers and Newfoundlands; PC3–4 resolve Bernese Mountain Dogs and Irish Wolfhounds.

UMAP visualization [McInnes et al., 2018] of the first 20 PCs (*n*_neighbors_ = 30, min_dist = 0.5) revealed discrete breed clusters (Figure 2). The “Labrador Or Golden Retriever” mixed group positioned intermediate between parent-breed clusters, providing biological validation. Coloring by source cohort confirmed the absence of systematic batch effects: breeds represented in multiple cohorts co-localized regardless of origin (Figure 2, right). The Wiener UK-origin Labrador Retrievers formed a sub-cluster distinct from US-origin Labradors, consistent with documented transatlantic population structure [Wiener et al., 2017]. ComBat batch correction [Johnson et al., 2007] was evaluated but rejected after investigation revealed that apparent batch effects were driven by cohort-breed confounding rather than technical artifacts.

### 4.3 Imputation quality

Imputation against the Dog10K reference panel yielded 9,667,790 variants passing filters (DR2 ≥ 0.3, MAF ≥ 0.01), a 125-fold densification over the typed scaffold (Figure 1C, Table 3). From 29.1M raw imputed variants, 23.2M passed DR2 filtering (79.7%), of which 9.7M also passed MAF filtering. The median perchromosome pass rate was 82.0%, with mean DR2 of 0.55–0.62 across non-problematic autosomes (Figure 3A– B). Ti/Tv ratios were consistent (2.1–2.7; Figure 3D).

**Figure 1:**
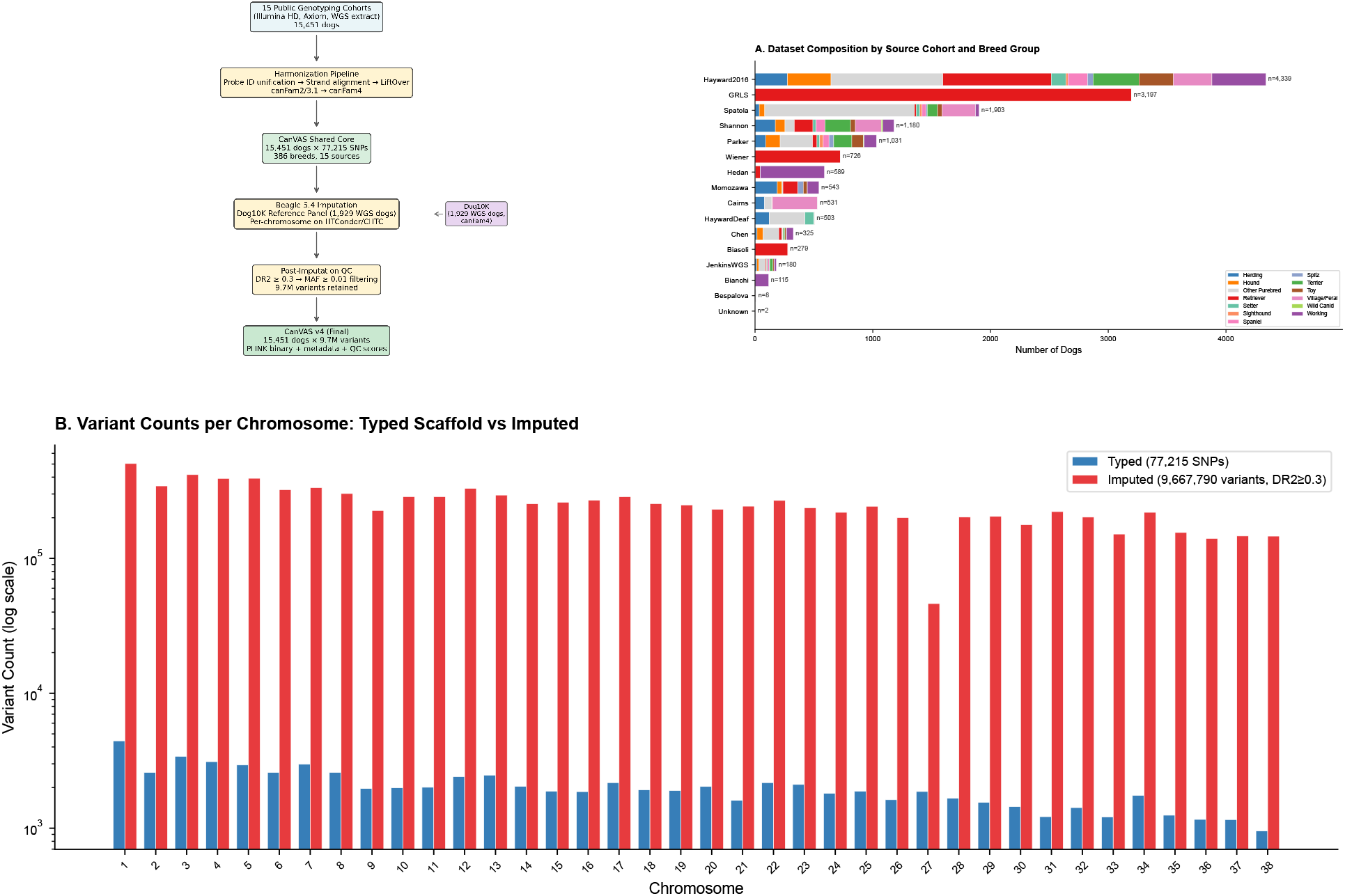
CanVAS dataset overview. Schematic of the harmonization and imputation pipeline, from 15 public genotyping cohorts through probe ID unification, strand alignment, and liftover to CanFam4, followed by Beagle 5.4 imputation against the Dog10K reference panel (1,929 whole-genome sequenced dogs). (A) Dataset composition by source cohort (rows) and breed group (colors), with sample counts indicated at right. Breed groups were assigned using AKC classifications. (B) Per-chromosome variant counts for the typed scaffold (77,215 SNPs; blue) and imputed dataset (9,667,790 variants passing DR2 ≥ 0.3 and MAF ≥ 0.01; red) on a log scale. The reduced imputed yield on chromosomes 27 and 32 reflects impaired imputation due to a documented coordinate reversal between the CanFam3.1 and CanFam4 assemblies.

**Figure 2:**
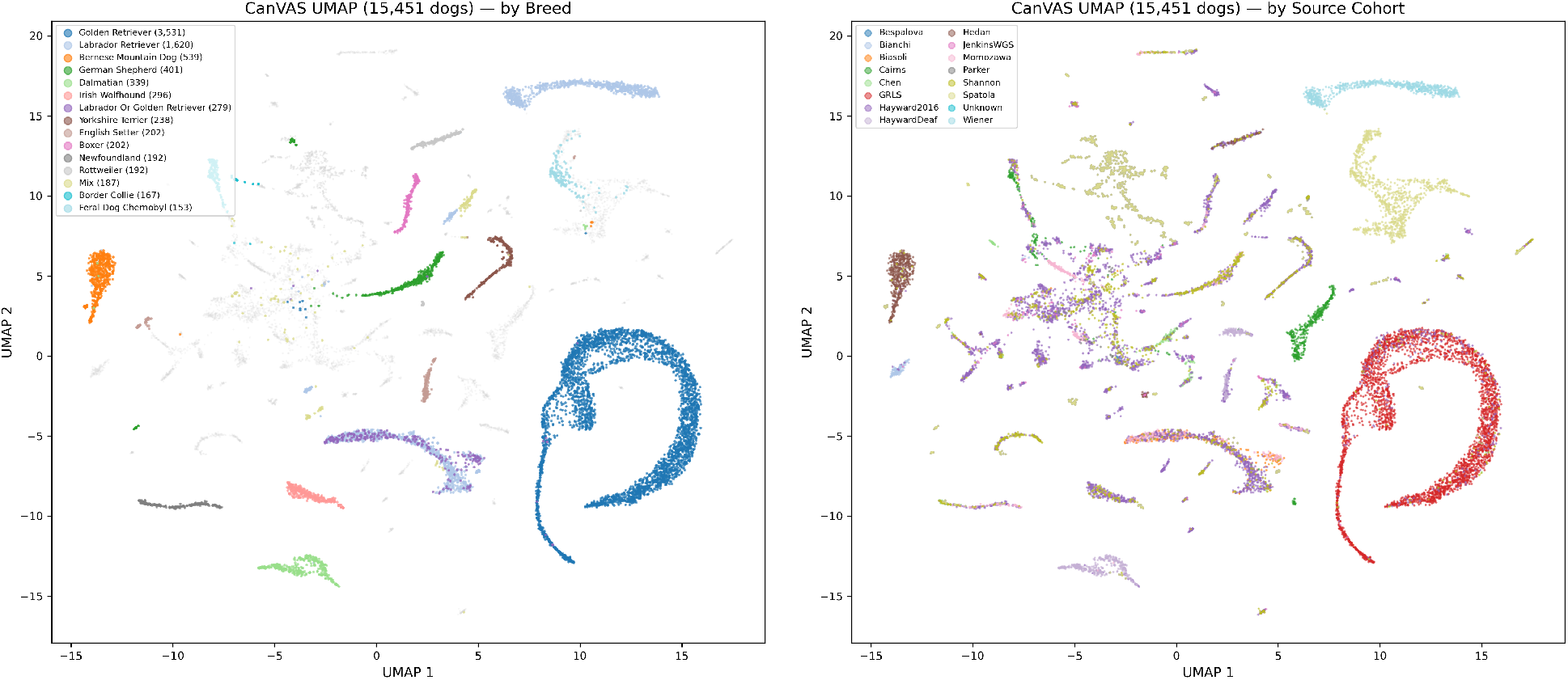
Population structure of 15,451 dogs via UMAP. Left: UMAP embedding colored by breed, with the 15 most abundant breeds labeled and sample sizes in parentheses; remaining breeds are shown in gray. Right: the same embedding colored by source cohort, demonstrating the absence of systematic batch effects; breeds represented in multiple cohorts co-localize regardless of origin. UMAP was computed on the first 20 principal components (*n*_neighbors_ = 30, min_dist = 0.5) from the LD-pruned typed backbone.

**Table 3:** Per-chromosome imputation summary. Chromosomes 27 and 32 (bold) show reduced quality due to the CanFam3.1/CanFam4 coordinate reversal.

| Chr | Total variants | Pass (DR2≥0.3) | High quality | Mean DR2 | Pass (%) |
| --- | --- | --- | --- | --- | --- |
| 1 | 1,555,491 | 1,267,697 | 525,526 | 0.60 | 81.5 |
| 2 | 1,089,269 | 883,515 | 340,717 | 0.59 | 81.1 |
| 3 | 1,212,717 | 1,002,561 | 447,382 | 0.62 | 82.7 |
| 4 | 1,156,131 | 949,768 | 428,078 | 0.61 | 82.2 |
| 5 | 1,189,352 | 964,416 | 391,262 | 0.59 | 81.1 |
| 6 | 987,568 | 799,927 | 333,565 | 0.60 | 81.0 |
| 7 | 994,434 | 817,666 | 368,005 | 0.61 | 82.2 |
| 8 | 936,287 | 757,544 | 301,520 | 0.59 | 80.9 |
| 9 | 733,598 | 574,408 | 189,964 | 0.55 | 78.3 |
| 10 | 889,786 | 716,070 | 270,248 | 0.58 | 80.5 |
| 11 | 891,389 | 726,733 | 256,257 | 0.58 | 81.5 |
| 12 | 954,659 | 788,266 | 337,891 | 0.61 | 82.6 |
| 13 | 864,559 | 712,648 | 309,956 | 0.61 | 82.4 |
| 14 | 765,890 | 621,371 | 263,945 | 0.60 | 81.1 |
| 15 | 796,004 | 647,243 | 258,519 | 0.59 | 81.3 |
| 16 | 809,853 | 638,981 | 226,089 | 0.56 | 78.9 |
| 17 | 847,688 | 698,319 | 292,507 | 0.61 | 82.4 |
| 18 | 774,637 | 622,741 | 256,273 | 0.59 | 80.4 |
| 19 | 743,134 | 610,218 | 249,874 | 0.60 | 82.1 |
| 20 | 715,796 | 580,251 | 230,889 | 0.59 | 81.1 |
| 21 | 683,206 | 557,853 | 219,986 | 0.59 | 81.7 |
| 22 | 806,091 | 658,499 | 281,149 | 0.60 | 81.7 |
| 23 | 674,121 | 556,681 | 245,820 | 0.61 | 82.6 |
| 24 | 646,665 | 529,467 | 201,934 | 0.59 | 81.9 |
| 25 | 711,241 | 581,977 | 231,181 | 0.60 | 81.8 |
| 26 | 547,877 | 459,338 | 180,495 | 0.61 | 83.8 |
| **27** | **587,054** | **62,238** | **4,481** | **0.08** | **10.6** |
| 28 | 587,363 | 483,093 | 180,585 | 0.59 | 82.2 |
| 29 | 577,429 | 477,494 | 190,511 | 0.60 | 82.7 |
| 30 | 504,007 | 414,476 | 155,080 | 0.60 | 82.2 |
| 31 | 619,646 | 516,515 | 176,288 | 0.59 | 83.4 |
| **32** | **519,495** | **272,308** | **3,882** | **0.29** | **52.4** |
| 33 | 438,486 | 360,825 | 127,284 | 0.59 | 82.3 |
| 34 | 622,622 | 517,767 | 214,953 | 0.61 | 83.2 |
| 35 | 407,973 | 344,919 | 130,772 | 0.61 | 84.5 |
| 36 | 414,375 | 347,363 | 116,600 | 0.59 | 83.8 |
| 37 | 436,992 | 361,474 | 128,131 | 0.59 | 82.7 |
| 38 | 396,930 | 333,080 | 105,153 | 0.58 | 83.9 |
| **All** | **29.1M** | **23.2M** | **8.4M** | **0.57** | **82.0** |

**Figure 3:**
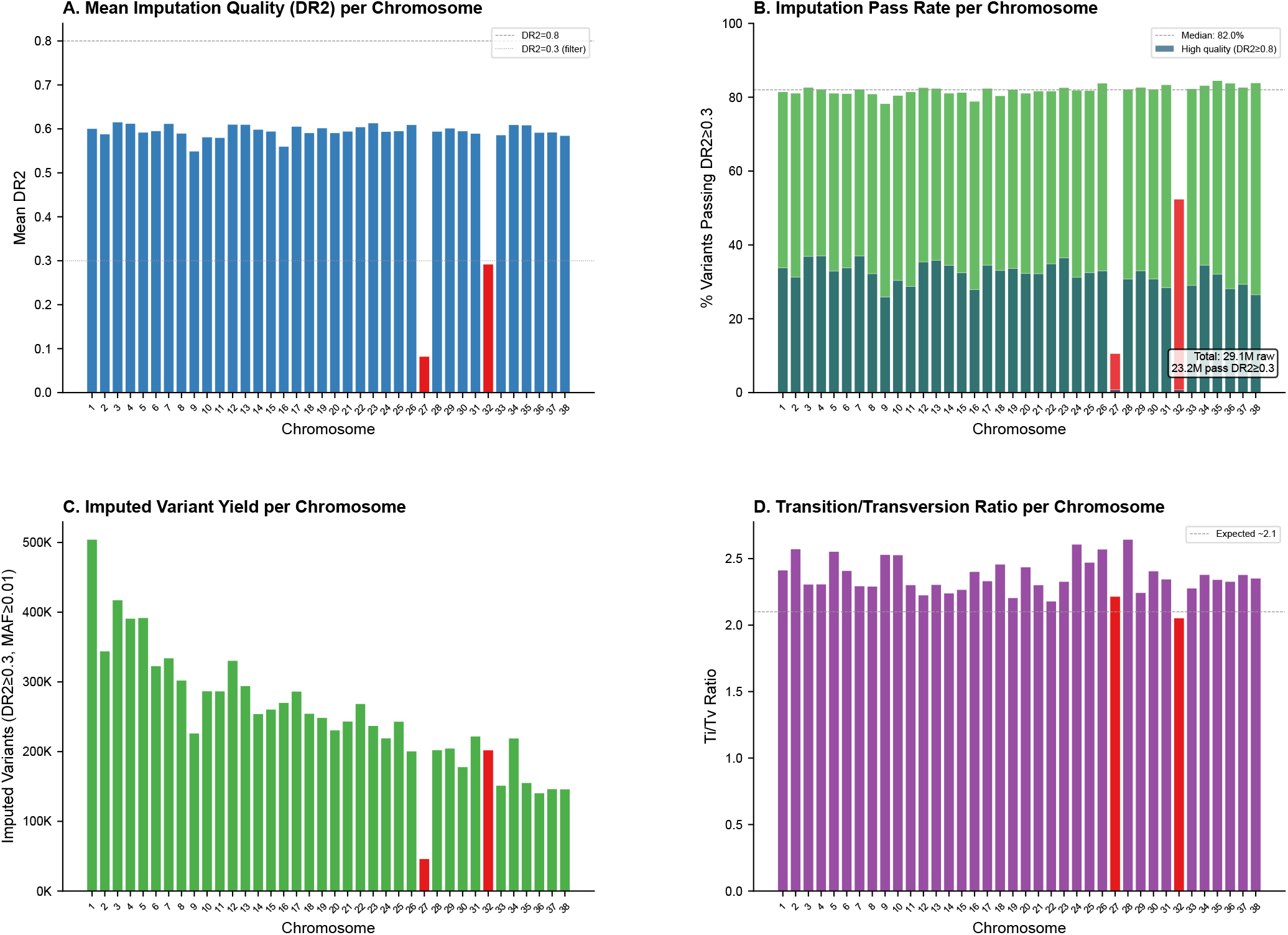
Imputation quality across the canine genome. (A) Mean imputation quality score (DR2) per chromosome. The upper dashed line indicates DR2 = 0.8 (high quality); the lower dashed line indicates DR2 = 0.3 (minimum filter threshold). Chromosomes 27 and 32 (red) show markedly reduced quality due to the CanFam3.1-to-CanFam4 coordinate reversal. (B) Percentage of variants passing the DR2 ≥ 0.3 filter per chromosome, with the high-quality subset (DR2 ≥ 0.8) shown separately. The dotted line indicates the genome-wide median pass rate (82.0%). (C) Total imputed variant yield per chromosome after DR2 and MAF filtering. (D) Transition-to-transversion (Ti/Tv) ratio per chromosome; the dashed line indicates the expected ratio of approximately 2.1.

Chromosomes 27 and 32 exhibited impaired imputation quality (mean DR2 = 0.08 and 0.29; pass rates 10.6% and 52.4%, respectively) due to the documented coordinate reversal between CanFam3.1 and CanFam4 [Wang et al., 2021, Field et al., 2020]. LD decay analysis confirmed that the typed backbone genotypes on these chromosomes are unaffected; only the imputation step is impaired.

The imputed dataset transformed the allele frequency spectrum (Figure 4). The typed backbone exhibited ascertainment bias toward common alleles (median MAF = 0.32; 495 variants with MAF *<* 0.05). Imputation recovered 2,977,396 rare variants (MAF *<* 0.05; 30.8%), yielding the L-shaped site frequency spectrum expected under neutral theory.

**Figure 4:**
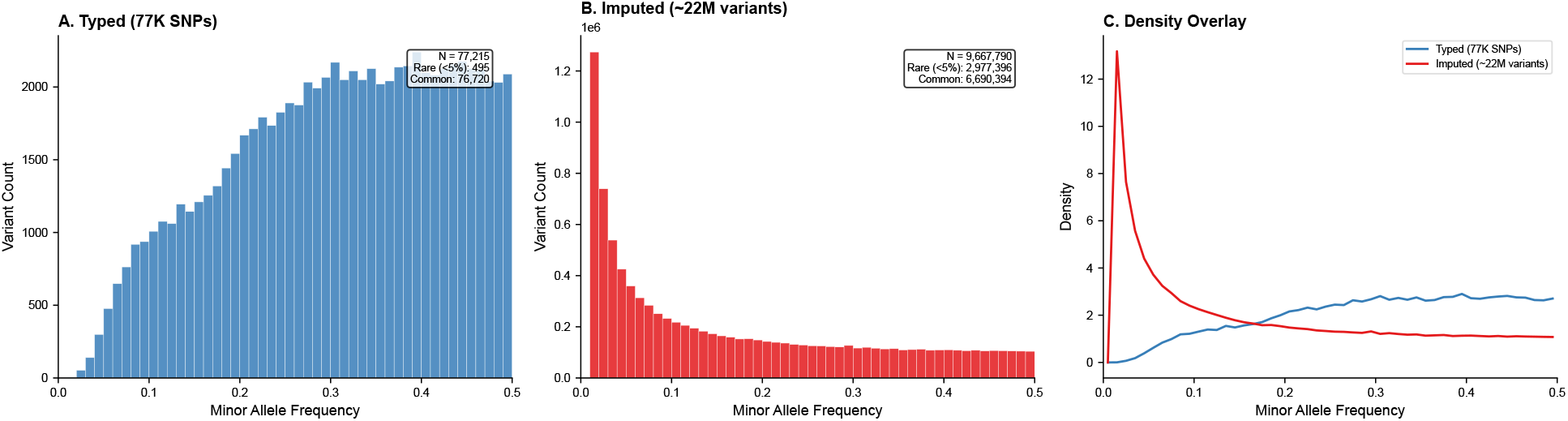
Minor allele frequency spectrum. (A) Site frequency spectrum of the typed scaffold (77,215 SNPs), showing ascertainment bias toward common alleles characteristic of SNP array platforms (median MAF = 0.32; 495 variants with MAF *<* 0.05). (B) Site frequency spectrum of the imputed dataset (9,667,790 variants), recovering the L-shaped distribution expected under neutral theory, including 2,977,396 rare variants (MAF *<* 0.05; 30.8%). (C) Density overlay comparing the typed and imputed frequency distributions.

### 4.4 Runs of homozygosity

To validate the resource’s utility for population genomics, we performed genome-wide ROH analysis on the typed backbone using PLINK (`--homozyg`; minimum 50 SNPs, minimum 500 kb, 1 heterozygote allowed per scanning window). The genomic inbreeding coefficient *F*_ROH_ was computed as total ROH length divided by autosomal genome size (2,410 Mb for CanFam4).

A total of 1,163,425 ROH segments were detected across 14,478 dogs, with mean *F*_ROH_ = 0.189. In-breeding varied widely: New Guinea Singing Dogs showed the highest values (median *F*_ROH_ = 0.63; *n* = 5), followed by Skye Terriers (0.54; *n* = 2) and Bull Terriers (0.49; *n* = 48), while village dogs (0.04–0.08) and mixed breeds (0.06) showed the lowest (Figure 5A–B). At the breed-group level, Setters and Working breeds exhibited the highest median *F*_ROH_; Village/Feral and Wild Canid groups the lowest (Figure 5B). ROH length class distributions varied across breed groups: Terrier and Setter breeds showed elevated long ROH (*>*8 Mb), while Retriever breeds showed predominantly medium-length ROH (2–8 Mb) (Figure 5C).

**Figure 5:**
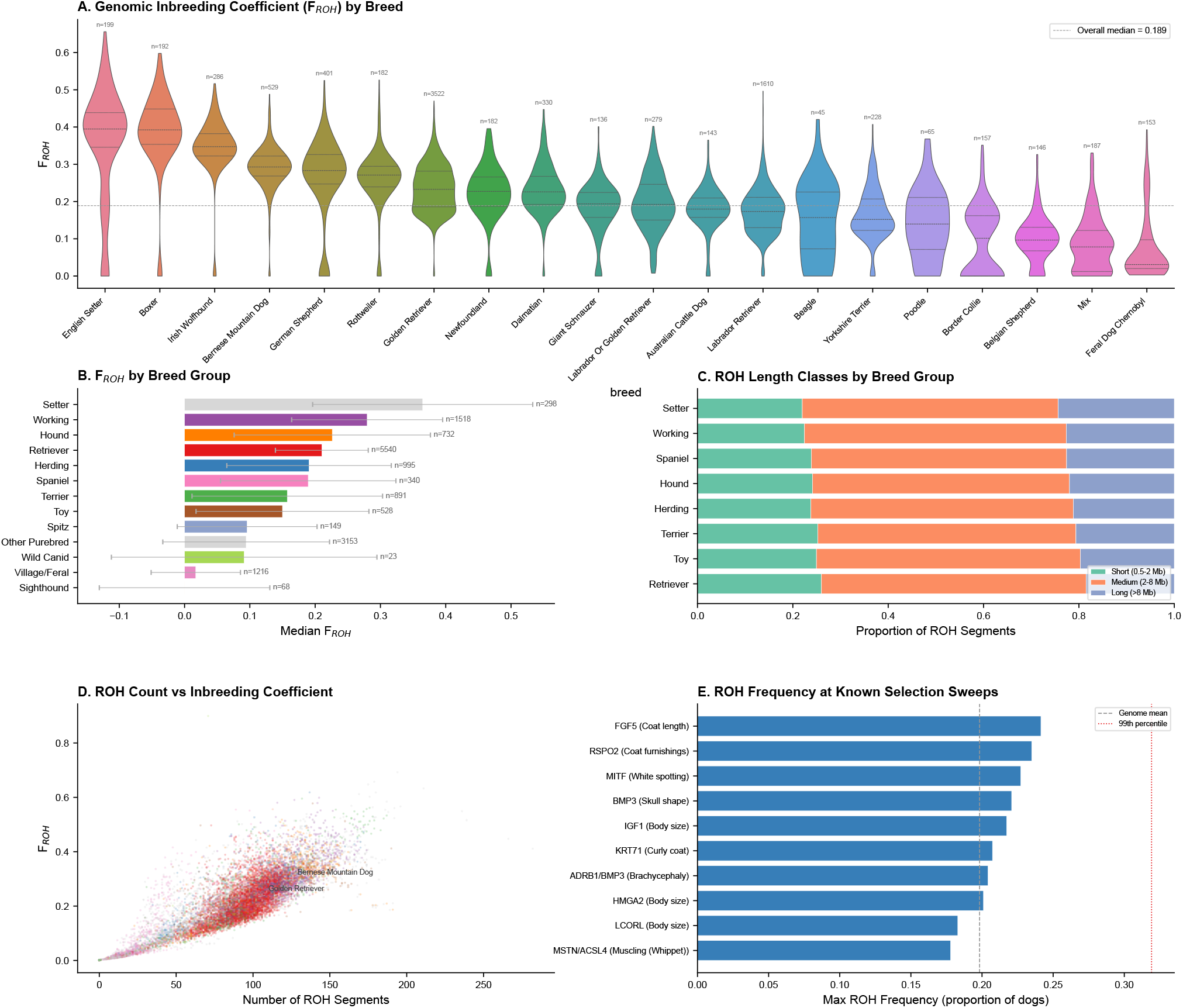
Breed-specific inbreeding landscape via runs of homozygosity. (A) Distribution of the genomic inbreeding coefficient (*F*_ROH_) for the 20 most abundant breeds (violin plots with embedded box plots; sample sizes above each violin). The dashed line indicates the overall median *F*_ROH_ (0.189). (B) Median *F*_ROH_ by breed group, with sample sizes at right and error bars showing interquartile range. (C) Proportion of ROH segments in three length classes: short (0.5–2 Mb, reflecting ancient shared ancestry), medium (2–8 Mb, consistent with historical bottlenecks or popular sire effects), and long (*>*8 Mb, indicating recent consanguinity). (D) Relationship between ROH segment count and *F*_ROH_ for individual dogs, with Golden Retrievers and Bernese Mountain Dogs highlighted. (E) Maximum ROH frequency at ten previously reported selection sweep loci, with the genome-wide mean (dashed) and 99th percentile (dotted) indicated for reference.

These patterns are consistent with previously reported breed-level differences in effective population size and breeding history.

ROH frequency at ten previously reported selection sweep loci ranged from 0.04 to 0.25, with none exceeding the genome-wide 99th percentile in the pooled multi-breed analysis, consistent with the breed-specific nature of these signals (Figure 5E).

### 4.5 Linkage disequilibrium

LD decay analysis confirmed long-range haplotype structure characteristic of the canine genome, with mean *r*^2^ declining from ∼ 0.65 at short distances to ∼ 0.10 by 500 kb. Chromosomes 27 and 32 showed comparable decay to other autosomes, confirming that typed backbone genotypes are unaffected by the assembly coordinate issue.

### 4.6 Known-locus validation

As a final check on the harmonization and imputation pipeline, we compared allele frequencies at four well-characterized trait loci between breed groups with known divergent phenotypes. At each locus, the most differentiated SNP showed clear frequency divergence in the expected direction in both the typed backbone and imputed dataset (Table 2). The stronger divergence in the imputed data reflects the higher variant density enabling identification of SNPs closer to causal variants, consistent with successful imputation of true biological signal.

## 5 Usage Notes

For the typed backbone, no imputation-related caveats apply. For the imputed dataset:

- Chromosomes 27 and 32 have reduced imputation quality and should be filtered or excluded for sensitivity analyses.
- Coordinates are on CanFam4; users working with CanFam3.1 annotations should liftover accordingly.
- DR2 scores are not retained in merged PLINK files.
- Cohort should be included as a covariate in association analyses.

## Data Availability

The CanVAS imputed genotype dataset, sample metadata, and quality control summary are available on Zenodo at https://doi.org/10.5281/zenodo.19186944 Brundage [2025]. The typed backbone PLINK files and all pipeline scripts are available at https://github.com/Brundage-VAIL/CanVAS. Per-chromosome VCF files with per-variant DR2 annotations can be regenerated from the deposited files using the provided pipeline scripts.

## Code Availability

All scripts are available at https://github.com/Brundage-VAIL/CanVAS under an MIT license. Key components include: `concordance.py` (allele concordance analysis), `liftover.py` (probe-ID-based coordinate remapping), `run_pipeline.sh` (end-to-end harmonization)

## Acknowledgments

We thank the investigators who generated and publicly deposited the source datasets that made this resource possible. We would like to thank Morris Animal Foundation, its staff members and all participants in the Golden Retriever Lifetime Study, including the dog owners, their golden retrievers and the Study veterinarians who made this work possible. Computational resources were provided by the UW–Madison Center for High Throughput Computing (CHTC).

## Author Contributions

D.M.B. conceived the study, developed the harmonization and imputation pipelines, performed all analyses, and wrote the manuscript.

